# vcfsim: flexible simulation of all-sites VCFs with missing data

**DOI:** 10.1101/2025.01.29.635540

**Authors:** Paimon Goulart, Kieran Samuk

**Affiliations:** Department of Computer Science and Engineering, The University of California, Riverside; Department of Evolution, Ecology, and Organismal Biology, The University of California, Riverside

**Keywords:** Software, Variant Call Format (VCF), Simulation, Coalescent, Population genetics, Missing data, Ploidy, Demography, Benchmarking, Genomic data analysis

## Abstract

**Background:** VCFs are the most widely used data format for encoding genetic variation. By design, standard VCFs do not include data from sites where all individuals are homozygous for the reference allele (“invariant sites”) and thus do not differentiate these from sites where data are completely missing. However, missing data are a key feature of biological datasets across all domains of genomics, and many recent studies have shown that missing data can introduce a variety of statistical biases in the estimation of key population genetic parameters. A solution to this limitation is to include invariant sites in a standard VCF, creating an “all-sites VCF”, exposing missing and invariant sites explicitly. One hurdle to the wider adoption of all-sites VCFs is a reliable parameterized simulation framework for generating biologically realistic all-sites VCFs.

**Results:** Here, we introduce an open-source command line tool, *vcfsim*, that interfaces with the popular coalescent simulation platform *msprime* and provides convenience functions for simulating all-sites VCFs with variable levels of ploidy and missing data. We show that the post-processed VCFs generated using *vcfsim* align precisely with population genetic expectations (i.e. are statistically identical to raw *msprime* output), accurately introduce missing data, and permit the simulation of data with varying ploidy levels, including the simulation of intraindividual ploidy variation (e.g. heterogametic sex chromosomes) and population structures.

**Conclusions:** Our results *vcfsim* is a useful and easy-to-use tool for the benchmarking of new software tools, performing population genetic inference, training of machine learning models, and the exploration of the effects of missing data in genomics data sets.

## Background

The Variant Call Format (VCF) is the most widely used file format for encoding genetic variation data, particularly in the context of population genetics and genomics studies (Danecek et al., 2011). VCF files are designed to store information about genetic variants, including single nucleotide polymorphisms (SNPs), insertions, deletions, and structural variants. Despite their widespread usage, VCFs have several key limitations. One significant drawback is that, by design, sites where all individuals are homozygous for the reference allele, i.e. invariant sites, are typically omitted (Danecek et al., 2011). This feature makes it impossible to distinguish between sites that are missing due to lack of sequencing data and sites that are simply invariant. This missing information complicates downstream analyses, particularly in cases where distinguishing between these types of sites is crucial, such as the estimation of nucleotide diversity within and between populations (Korunes & Samuk, 2021; Samuk, 2023).

One extremely common workaround is the use of “all-sites” VCFs, which include both variant and invariant sites. By explicitly including invariant sites, missing data are exposed – missing sites are truly missing and not either missing or invariant. All modern variant calling pipelines (GATK, bcftools, freebayes, etc., cite) include the ability to produce all-sites VCFs. Popular tools such as *pixy* (Korunes & Samuk, 2021) and piawkwa (cite) make use of all-sites VCFs to calculate population genetics statistics while accounting for missing data and per-site differences in sample size. However, benchmarking tools that make use of all-sites VCFs has been challenging due to the lack of native support from common simulation packages, which typically output standard (i.e. variants only) VCFs. Simulating all-sites VCFs typically involves complex and ad-hoc workflows, especially when missing data and other biological details are involved.

Indeed, missing data are widely known to introduce serious biases in the estimation of a variety of population genetic parameters (Korunes & Samuk, 2021; Lowry, 2017; Samuk, 2023; Wong, Zeng, & Lin, 2019), yet there are no widely adopted tools for simulating ground-truth data with predefined levels of missing genotypes or sites. This is an issue, as researchers often need simulated data with specific patterns of missing information or population genetic parameters to validate tools or analyses. However, current methods require ad-hoc solutions and lack flexibility. As such, there is a need for standardized tools to simulate population genetic data and output all-sites VCFs, both for working population genetics researchers and bioinformaticians developing new tools.

Here, we introduce *vcfsim*, a command-line tool that provides a user-friendly interface that produces all-sites VCFs using the coalescent simulation package *msprime* (Baumdicker et al., 2022) with customizable levels of missing data. The tool also supports simulation of VCFs across various ploidy levels, including mixed ploidies (e.g., diploid organisms with haploid sex chromosomes), as well as simulation of multiple evolving populations. This makes *vcfsim* a flexible and accurate tools for rapidly simulating genetic data. By addressing the limitations of current tools, *vcfsim* simplifies the process of creating biologically realistic VCFs for use in testing and validating genetic analysis pipelines, as well as in inferential tasks needing large amounts of simulated data such as approximate Bayesian computation.

## Methods

We developed *vcfsim* in Python version 3.11 leveraging tools from the following libraries: ipython (Perez and Granger 2007), numpy (Harris et al. 2020), and pandas (The pandas development team 2024). The coalescent simulation package msprime (Baumdicker et al., 2022) provides the core population genetic-based backend for the simulation. The source code for *vcfsim* can be found at http://github.com/samuk-lab/vcfsim, and is freely available via MIT license. It can also be easily installed via the package management platform conda (conda contributors 2025), on bioconda (Grüning et al. 2018): https://bioconda.github.io/recipes/vcfsim/README.html

*vcfsim* simulates realistic genetic variation data that can be highly customized to match a wide variety of use cases. Our focus was on a easy to deploy, one-step solution for simulating VCFs for all kinds of biological applications. Unlike tools such as *vcfgl* (Altinkaya, 2025), our tool was designed for hard-filtered VCF workflows (i.e. not operating on genotype likelihoods), which are extremely common in most fields. Our tool *vcfsim* works by simulating genetic data from a single population using a neutral coalescent process (via *msprime*), applying mutations to these sequences, outputting the results in “all-sites” VCF format, and post-processing this VCF to simulate missing data of various kinds and variable ploidy. Users can customize the simulation via command-line arguments to specify the desired characteristics of the simulated population and VCF output. Key features of *vcfsim* include:

- Random seed control: Users can specify a random seed to control stochastic elements of the coalescent simulation and missing data processes. Identical seeds yield identical results for repeated runs, ensuring reproducibility.
- Missing data simulation: The software allows for the simulation of missing data, both at the level of genotypes and whole sites.
- Flexible chromosome inputs: Users can simulate multiple chromosomes simultaneously by providing an input file with parameters for each chromosome (e.g. multiple autosomes and a sex chromosome).
- Support for arbitrary ploidy: Any ploidy level supported by msprime can be simulated, which we extended to handle mixed ploidies such as those found in systems with heterogametic sex chromosomes
- Multiple populations: vcfsim can simulate samples drawn from two populations with a shared coalescent history, with varying divergence times and population sizes.
- Efficient handling of large simulations: *vcfsim* supports running multiple simulation replicates from a single command, facilitating use for tool validation, statistical model fitting, and machine learning model training.

### Parameters and inputs

*vcfsim* offers a range of user-specified customization options. These parameters include both required and optional arguments that control various aspects of the simulation process, such as the random seed for reproducibility, the level of missing data, and the characteristics of the population being simulated. The full list of required and optional parameters and given in Table 1 and Table 2 respectively.

**Table 1.**
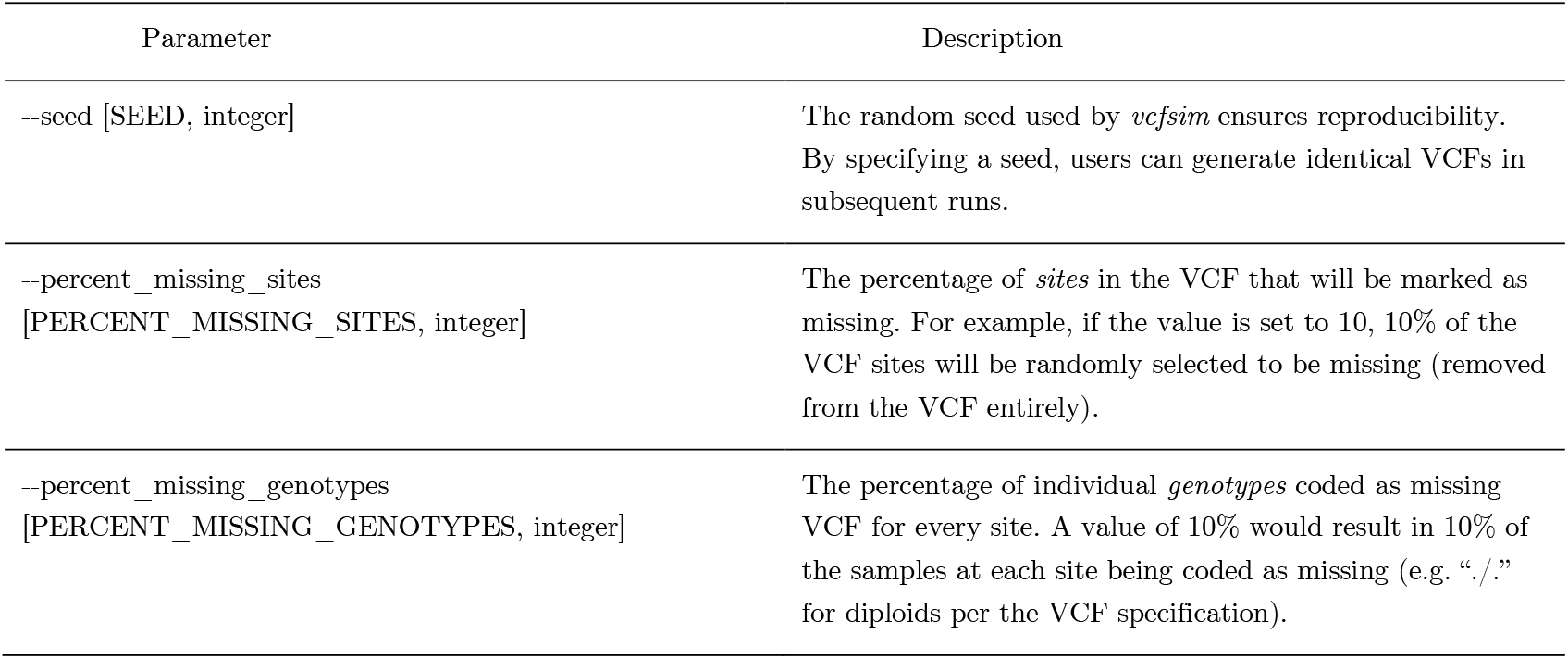
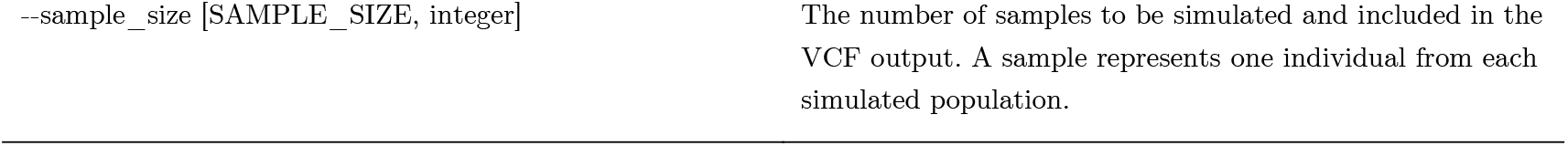
Required parameters for vcfsim. Each row represents a single required parameter, with its name, value (in capital letters), and type of parameter (e.g. an integer, string, etc.). Square brackets indicate placeholders for actual parameter values. The description column contains a description of the function of each parameter in controlling the behavior of vcfsim.

**Table 2.**
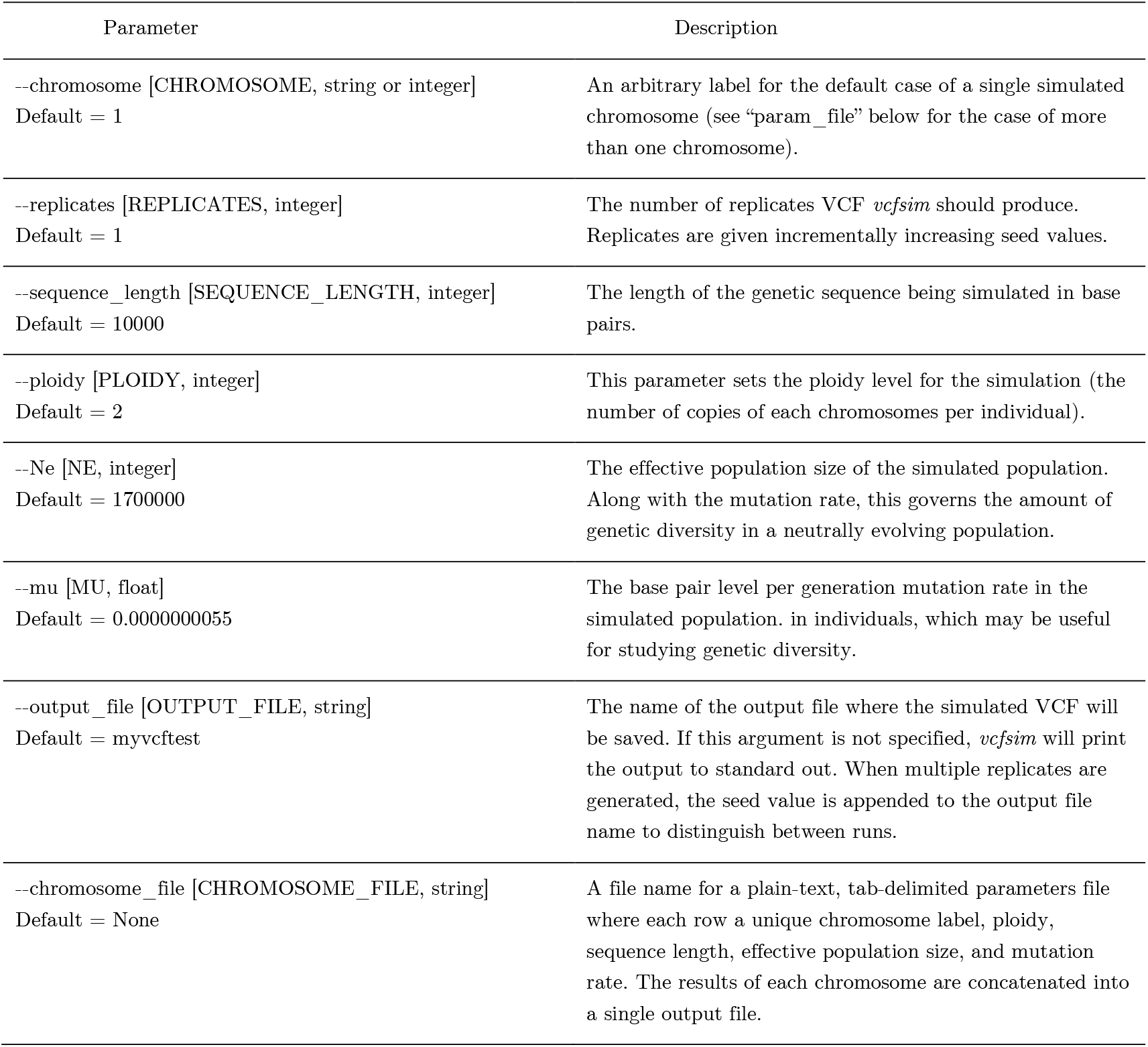
Optional parameters for vcfsim (i.e. those not required for a run). Each row represents a single required parameter, with its name, value (in capital letters), and type of parameter (e.g. an integer, string, etc.). Square brackets indicate placeholders for actual parameter values. The description column contains a description of the function of each parameter in controlling the behavior of vcfsim.

### Output options

Users can specify an output file for the generated VCF data using the “--output_file” argument. The resulting file will always be in the .vcf format and will also append the random seed used for the simulation at the end of the filename ensuring repeatability of results. The exact command line parameters invoked to produce a specific VCF are also encoded in the “source” metadata field of each VCF. If no output file is specified, the VCF data will be printed directly to standard output. This feature allows users to integrate *vcfsim* easily into command line pipelines, enabling further processing or analysis without the need to generate a static file.

### Validation of accuracy

To assess the biological and numerical accuracy of *vcfsim*, we performed a variety of validation procedures. The accuracy of *msprime* has been thoroughly demonstrated (Baumdicker et al., 2022), however *vcfsim* performs substantial post-processing and manipulation of VCFs produced by *msprime* via the addition of invariant sites and missing data, multiple chromosomes, and variable ploidy. As such, we sought to validate the following:

First, to evaluate whether the simulated VCFs generated by *vcfsim* align with theoretical expectations of nucleotide diversity (π), we generated 10,000 diploid VCF files, each with a mutation rate of 1e-9, an effective population size (N_e_) of 1,700,000, a sequence length of 10 000 base pairs, and a sample size of 10 (i.e. 10 diploid genotypes). Each simulation was performed with a unique random seed, starting from 1 000 and incrementing by 1 up to 11 000. To measure π for each simulated VCF, we employed both VCFtools (Danecek et al., 2011) and pixy (Korunes & Samuk, 2021) to ensure consistency between the tools and to verify that *vcfsim* did not introduce any errors. This dual approach also allowed us to cross-check the π values calculated by both libraries.

To verify that the percent of missing genotypes and missing sites introduced by *vcfsim* align with the specified input values, we obtained an independent validation of the extent of missing data using *bcftools* (Danecek et al., 2021). For missing genotypes, we generated VCF files with a sample size of 100 and incrementally introduced missing genotypes, ranging from 0% to 100%, increasing by 1% (one missing sample) with each iteration. Similarly, to assess the accuracy of missing site generation, we created another script to manually count the number of missing rows in the VCF files. In this case, we set the sequence length to 100 and progressively introduced missing sites, again in 1% (one site) increments from 0% to 100%. Ideally, we expect a perfect linear relationship between the specified and measured values, confirming that *vcfsim* is accurately applying the desired levels of missing data.

To assess runtime and assess overall computational efficiency, we used *vcfsim* to simulate sets of 1000 VCFs with the following parameters: 10 diploid samples, mutation rate=1e-9, N_e_=1.7M, and sizes of 1 000-1 000 000bp, in steps of powers of 10.

Finally, we provide two worked examples of the generation of datasets with variable ploidy: (1) a tetraploid organism, and (2) a diploid organism with two autosomes and a pair of heterogametic sex chromosomes (e.g. similar to a XY human male). In accordance with Wilson Sayres (2018), we again computed π, with the expectation of the following levels of average genetic diversity: autosomes π = 4N_e_μ, X chromosome π = 3N_e_μ (¾ of autosomes), and Y chromosome π = N_e_μ (¼ of autosomes). We used the following parameter values for these examples: N_e_ = 1M, μ = 1e-8, sequence length = 10000, and sample size = 10. All code used in the validation procedures and example plots for *vcfsim* are available as a Github repository: https://github.com/samuk-lab/vcfsim-benchmarking-and-test-analysis.

## Results

### Accuracy

The estimated π values calculated from *vcfsim* VCFs using both pixy and VCFtools were identical and closely matched theoretical expectations (Figure 2A, Figure 2B; 4Neμ = 0.0068; VCFtools mean π = 0.00676, 95% CI 0.00669–0.00682; pixy mean π = 0.00676, 95% CI 0.00669– 0.00682, n = 10000). We also observed a perfect linear relationship between the specified amounts of missing data specified by *vcfsim* and the values for missing data measured using *bcftools* and/or the raw count of rows in the VCF (Figure 2C, Figure 2D; R^2^ = 1.00, correlation test: p < 0.00001, n = 100)). This indicates that *vcfsim* correctly implements arbitrary levels of missing genotypes and sites.

**Figure 1.**
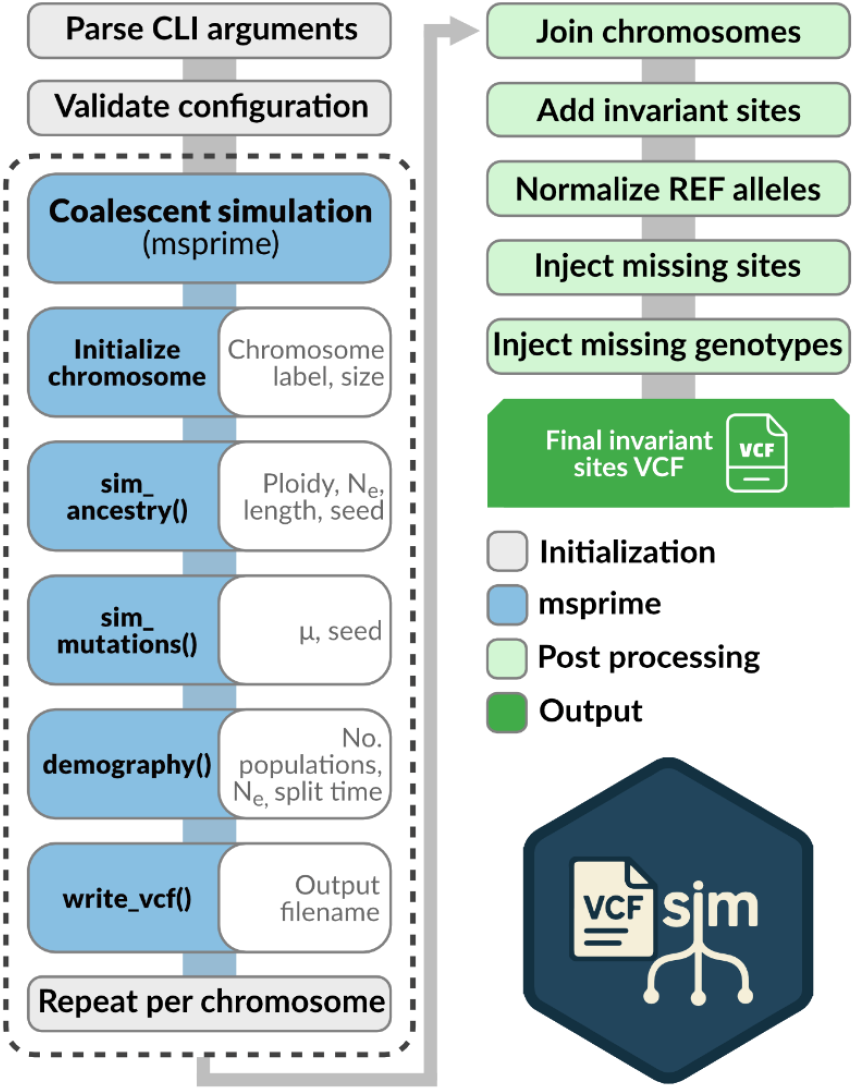
Flow chart showing the basic operation of *vcfsim*. Each step in the software (bubbles) is coded by its general function (see inset legend). The general flow of data through the program is indicated by the dark gray/blue line behind the steps. The coalescent simulation portion (dashed outlined) is repeated for each chromosome in multichromosome mode. The *msprime* simulation steps have user specified parameters (white interior bubbles) parsed from the command line invocation of *vcfsim*.

**Figure 2.**
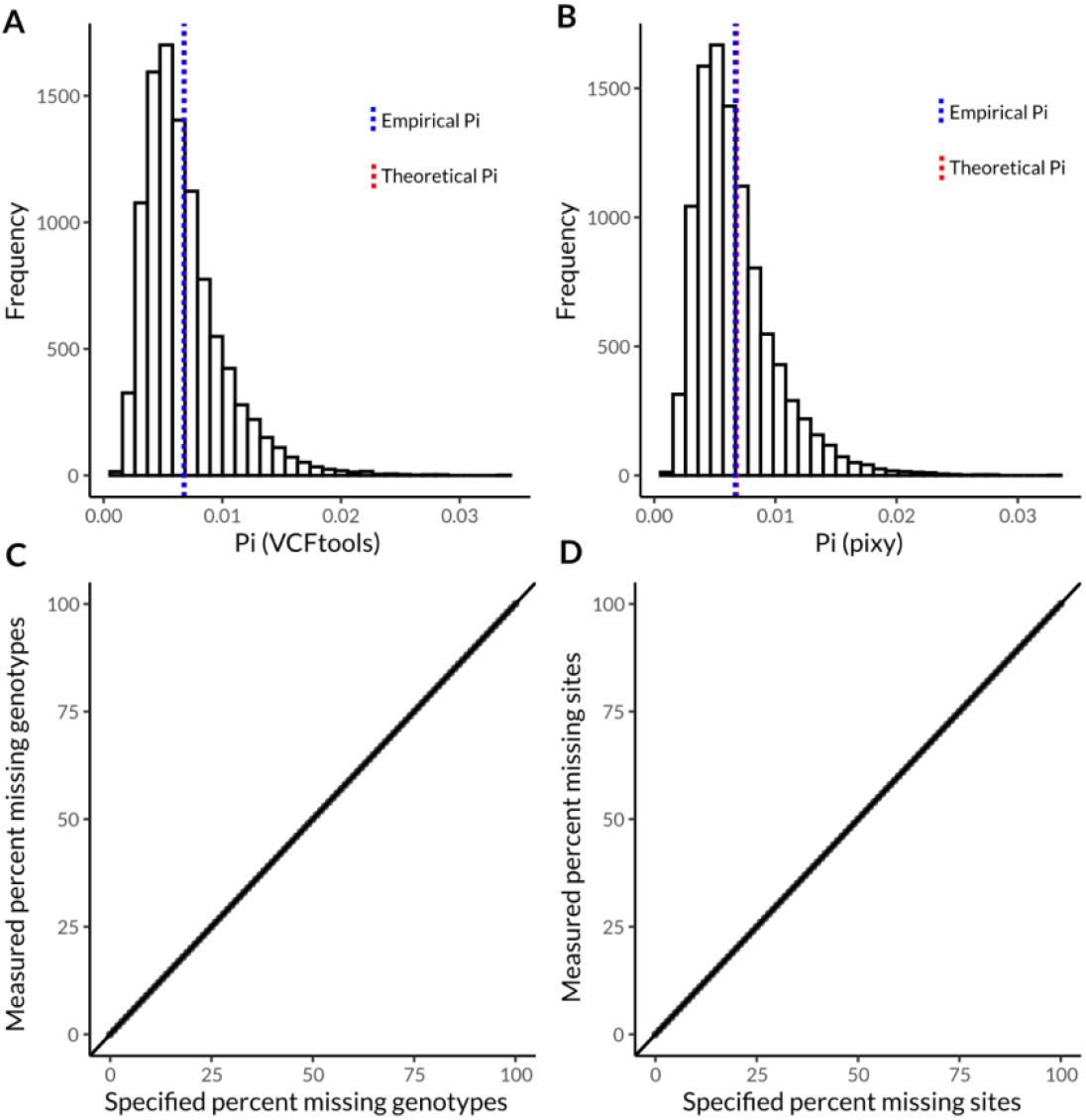
Distribution and accuracy of nucleotide diversity (pi) values and missing data in VCFs produced by *vcfsim*. (a) Distribution of pi values calculated with VCFtools in the absence of missing data. (b) Distribution of pi values calculated with pixy in the absence of missing data. Vertical lines in (a) and (b) represent the theoretical expectation for pi (red, 4N_e_μ), and the empirical mean of the computed values (blue). (c) Relationship between specified (*vcfsim*) and measured (bcftools) percentages of missing genotypes. (d) Relationship between specified (*vcfsim*) and measured (manual count) percentages of missing sites.

### Runtime

The runtime to create VCFs (10 diploid samples, mutation rate= 1e-9, Ne=1.7M) ranged from an average 0.649 seconds for a 1kbp VCF to 168 seconds for a 1Mbp VCF (Figure 3, means derived from 1000 replicate simulations). Using vcfsim, this task shows an approximate scaling relationship of ∼O(N^0.8^) complexity, i.e. efficient and potentially sublinear (Figure 3, Best fit power law exponent: T∝N^0.815^, 95% CI for k=0.466, 1.164).

**Figure 3.**
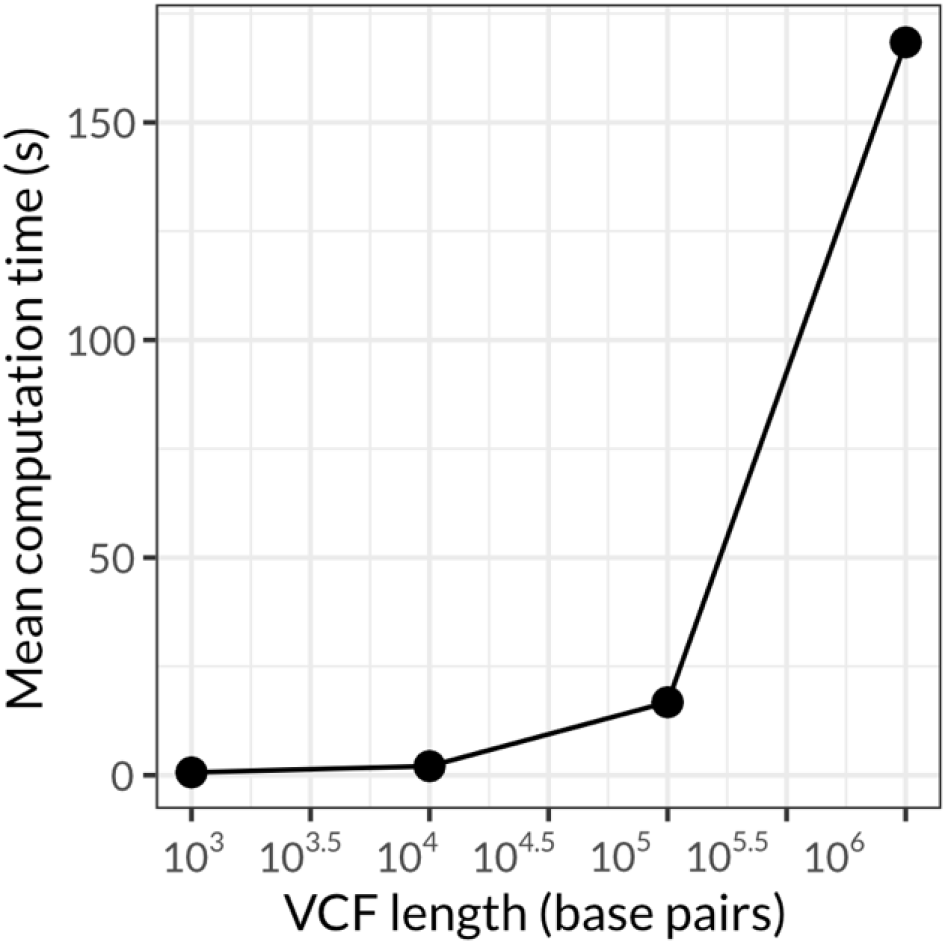
Mean computation time in seconds of VCFs of varying base pair length by *vcfsim*. Each point depicts the mean computation time of 1000 VCFs of the specified length (see “Runtime” Results section for full *vcfsim* parameterization). VCF length is shown on a log_10_ scale.

### Worked examples

To showcase additional features in *vcfsim*, we generated representative diploid VCFs displaying missing data (genotypes and sites, Figure 4A), data from a tetraploid organism (Figure 2B), as well as data from a diploid organism with heterogametic sex chromosomes (Figure 5). When multiple ploidy levels are simulated in this manner, the genetic diversity of autosomes and sex chromosomes are consistent with theoretical expectations (N_e_ = 1 000 00, μ = 1e-8; Figure 3, 4N_e_μ = 0.04, for autosomes, 3N_e_μ = 0.03 for X chromosomes, and N_e_μ = 0.01 for the Y chromosome).

**Figure 4.**
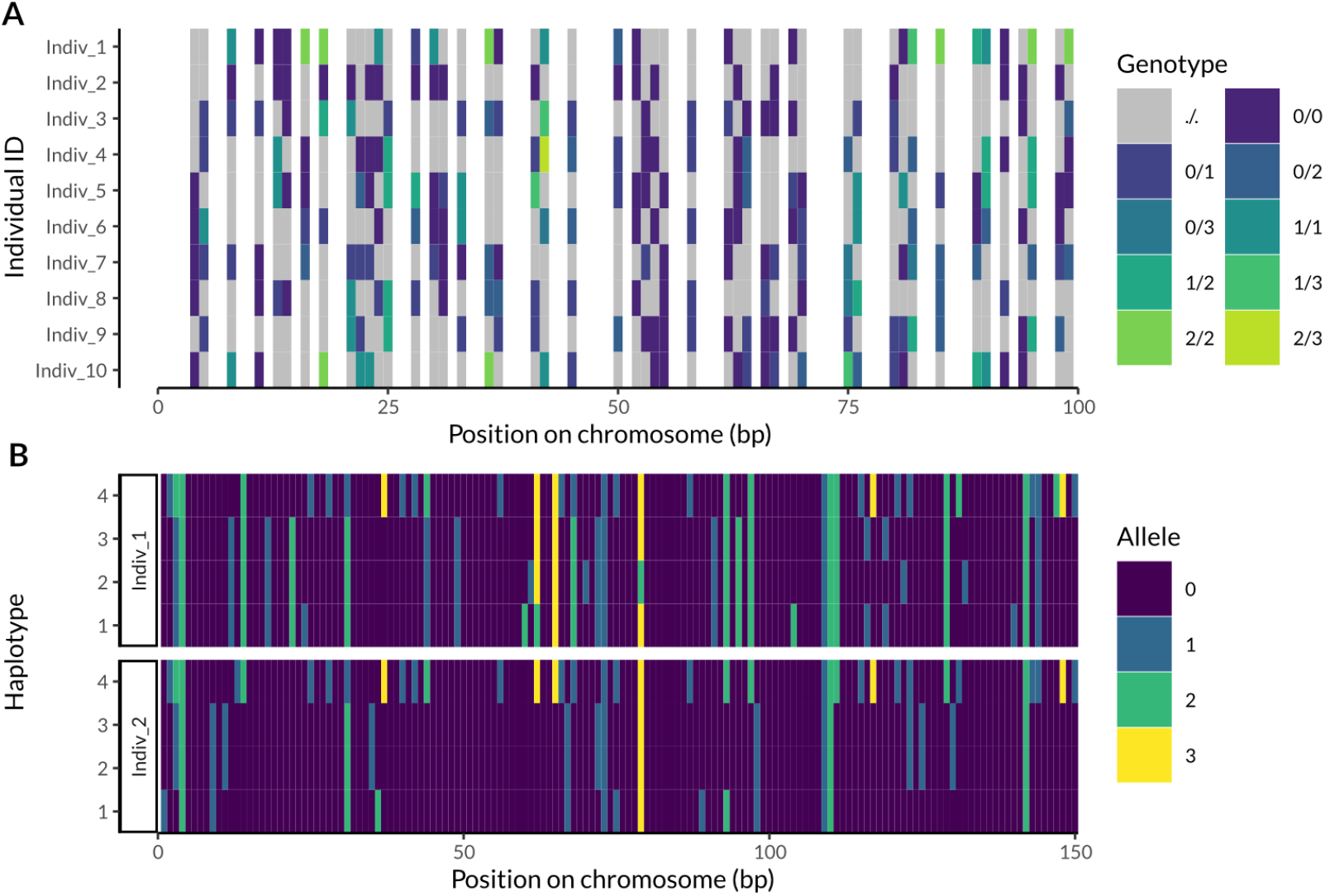
A heatmap visualization of VCF data generated using *vcfsim*. (A) A 100 bp span of genetic data from ten diploid individuals simulated using *vcfsim*. Each column represents a site in the genome, and each row a diploid individual. Each cell is colored based on its diploid genotype (note some sites are polyallelic), with grey representing a missing genotype. Completely missing sites are represented as white. (B) Simulated VCF of data from two tetraploid individuals with no missing data. Each column represents a site in the genome, and each row one of the individual haplotypes (1-4) for each individual (Indiv_1 and Indiv_2), with cells colored based on the haploid genotype (i.e. allele) at each site. Alleles in both plots are arbitrarily coded 0-4, with zero being the reference allele.

**Figure 5.**
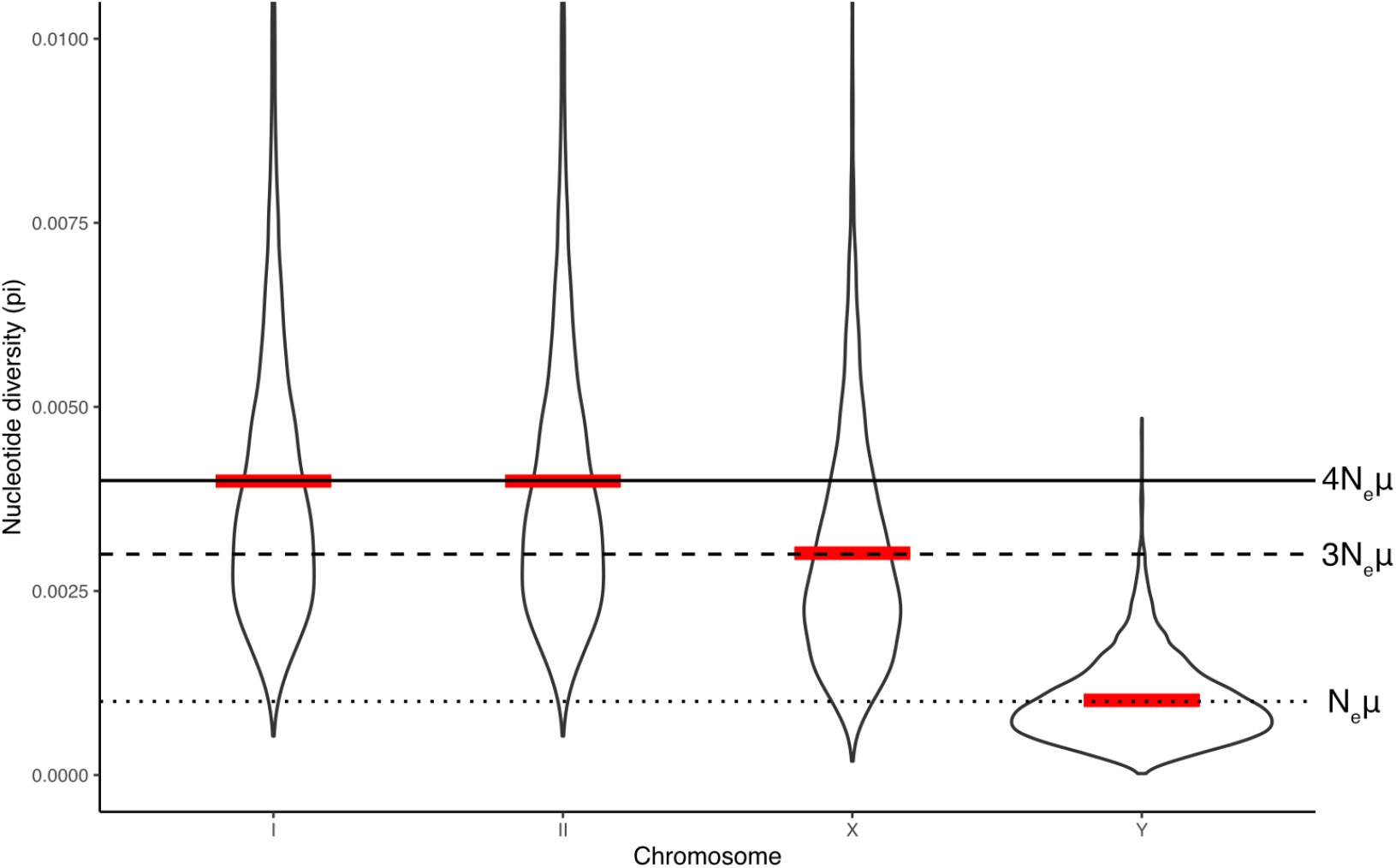
A violin plot of nucleotide diversity (pi) measured in 10000 bp windows in a diploid individual with autosomes (I, II) and heterogametic sex chromosomes (X, Y). Each violin shows the frequency distribution (density) for nucleotide diversity on each chromosome. Horizontal lines show the theoretical expectation for each class of chromosome (autosomes = 4N_e_μ, X = 3 N_e_μ, Y = N_e_μ). Red lines depict the empirical mean nucleotide diversity for the whole chromosome (expected to match the horizontal black lines).

## Discussion

*vcfsim* provides a standardized method for generating all-sites VCFs, complete with configurable levels of missing data and flexible ploidy and chromosome configurations. It provides a simple and customizable interface to the widely used coalescent simulation msprime (Baumdicker et al., 2022), along with parameterized postprocessing of VCFs to add invariant and missing sites. The ability to customize these parameters allows for the creation of test datasets that closely mimic real-world sequencing outputs, making it a useful tool for a variety of applications in genomics and population genetics.

We demonstrate that the VCFs generated by *vcfsim* align with the theoretical expectations of nucleotide diversity, with no errors introduced by the simulation or post-processing steps (Figure 2 C-D). Both tools we employed for computing nucleotide diversity (vcftools, pixy) show identical results (Figure 2A-B). Missing data (both sites and individual genotypes) are also exactly represented as specified in all output files. Together, these results validate the accuracy of *vcfsim* in replicating expected genetic diversity and missing data in simulated VCF files.

Our results also demonstrate that *vcfsim* is highly flexible, and capable of generating VCFs with varied ploidy levels and missing data. For example, simulations of tetraploid organisms (Figure 4B) and diploid organisms with heterogametic sex chromosomes (Figure 5C) accurately reflect theoretical diversity expectations. As such, *vcfsim* will be a useful tool across a wide variety of biological systems.

While a relatively straightforward tool, *vcfsim* fills an important gap in available tools and has many potential applications. One use of *vcfsim* is to benchmark software tools designed for genetic analysis, ensuring that they perform correctly across different configurations of missing data or ploidy. Additionally, *vcfsim* could be used to generate datasets for training different machine learning models, such as neural networks aimed at identifying genetic variants or inferring population structure. These datasets could also be used to conduct power analyses for population genetic inference, helping researchers assess how different levels of missing data or sample sizes affect the ability to detect evolutionary signals or demographic events.

### Limitations

While *vcfsim* provides an accurate and flexible tool for simulating VCFs, there are several important limitations to consider. First, *vcfsim* simulates VCFs with either completely known or completely missing data. In real-world sequencing data, however, we often encounter intermediate cases, such as sites with low sequencing depth or uncertain genotype likelihoods. *vcfsim* does not currently simulate these complexities (but see *vcfgl*, Altinkaya 2025). Another limitation is the size of the all-sites VCFs generated by *vcfsim:* because these files include both variant and invariant sites, they can become quite large, particularly for larger genomes. While the inclusion of invariant sites is essential for certain types of analyses (Korunes & Samuk, 2021), it can create challenges in terms of storage and computational efficiency (which are also true for empirical, non-simulated all-sites VCFs). Handling large all-sites VCFs may require significant memory and processing resources, which can be a bottleneck in large-scale simulations or analyses.

## Conclusion

*vcfsim* provides a simple, one-step solution for generating all-sites VCFs with customizable missing data and ploidy configurations. By addressing several limitations in current VCF generation workflows, *vcfsim* simplifies the process of creating realistic test datasets for use in genomic research and software validation. The ability to accurately simulate and format genetic variation and introduce controlled amounts of missing data opens up new possibilities for benchmarking tools, training machine learning models, and conducting power analyses in population genetics.

## Data Availability

The source code for vcfsim is available under an MIT license here: http://github.com/samuk-lab/vcfsim. The code used to perform benchmarking and accuracy validation is available under an MIT license here: https://github.com/samuk-lab/vcfsim-benchmarking-and-test-analysis.

## Declarations

Salary expenditures and compute time resources (1% of the project total of $1.25M) for the research reported in this publication was supported by NIGMS of the National Institutes of Health under award number 1R35GM154837-01 (MIRA). The content is solely the responsibility of the authors and does not necessarily represent the official views of the National Institutes of Health.

## Notes

https://github.com/samuk-lab/vcfsim

### Competing Interest Statement

The authors have declared no competing interest.

### Summary of Updates

We performed more extensive benchmarking, updated the software to allow for the simulation of multiple populations.

